# Aurora Kinase A Inhibition plus Tumor Treating Fields Suppress Glioma Cell Proliferation in a Cilium-Independent Manner

**DOI:** 10.1101/2023.11.29.569194

**Authors:** Jia Tian, Julianne C. Mallinger, Ping Shi, Dahao Ling, Loic P. Deleyrolle, Min Lin, Habibeh Khoshbouei, Matthew R. Sarkisian

## Abstract

Tumor Treating Fields (TTFields) have been shown to extend the survival of glioblastoma (GBM) patients. TTFields interfere with a broad range of cellular processes which may contribute to their efficacy. Among these, TTFields disrupt primary cilia stability on GBM cells. Here we asked if concomitant treatment of TTFields with other agents that interfere with GBM ciliogenesis can further suppress GBM cell proliferation in vitro. Aurora Kinase A (AURKA) promotes both cilia disassembly and GBM growth in vitro and in xenograft models. Inhibitors of AURKA such as Alisertib have been previously demonstrated to inhibit cilia disassembly and increase the frequency of cilia in various cell types. However, here we show that physiological concentrations of Alisertib treatment significantly reduced GBM cilia frequency in gliomaspheres across multiple patient derived cell lines, and in patient biopsies treated ex vivo with Alisertib. This activity of Alisertib seems to be glioma cell specific as it did not reduce neuronal or glial cilia frequencies in mixed primary cell cultures from mouse forebrain. Furthermore, Alisertib depletion of glioma cilia appears specific to AURKA inhibition, as a potent AURKB inhibitor, AZD1152, had no effect on GBM ciliary frequency. Treatment of two different GBM patient-derived cell lines with TTFields and Alisertib resulted in a significant reduction in cell proliferation compared to either treatment alone. However, this effect was not cilia-dependent as the combined treatment reduced proliferation in cilia-depleted cell lines lacking, *ARL13b*, or U87MG cells which are naturally devoid of ARL13B^+^ cilia. This result is not surprising given the wide range of pathways regulated by AURKA in addition to cilia. Nonetheless, Alisertib-mediated effects on glioma cilia may be a useful biomarker of drug efficacy within tumor tissue. Considering Alisertib has been shown to cross the blood brain barrier and inhibit intracranial growth of xenografted tumor models, our data warrant future studies to explore whether concomitant Alisertib and TTFields exposure prolongs survival of brain tumor-bearing animals in vivo.

## Introduction

Tumor Treating Fields (TTFields) therapy is the latest approved therapeutic option for glioblastoma (GBM). TTFields are low intensity (1-3V/cm), intermediate frequency (200kHz) alternating electric fields applied across the tumor through a wearable headset array. When combined with standard of care chemotherapeutic temozolomide (TMZ), overall GBM patient survival is prolonged by 4-5 months(1). How this synergism works remains unclear because TTFields engages a wide range of anti-tumor mechanisms on cells (2). Recently we reported that one cellular target affected by TTFields is the primary cilium of tumor cells in vitro and ex vivo (3). Primary cilia are microtubule-based organelles that are gaining increasing attention on GBM cells (4, 5). For example, cilia are linked to glioma cell stemness (6), angiogenesis (7) and chemoresistance (3, 8-10). In addition, our group and others have found that TMZ promotes the formation of primary cilia on GBM cells(3, 9, 10). Ablating glioma cilia genetically or with TTFields lowered resistance to TMZ (3, 8-10). This led us to explore whether targeting other tumor cell pathways essential for ciliogenesis could further enhance the effects of TTFields.

One critical regulator of ciliogenesis is AURKA (11), a molecule that has been extensively characterized in promoting GBM growth and treatment resistance. AURKA is frequently overexpressed in GBM compared to control and other brain tumor subtypes, and an inhibitor of AURKA, Alisertib (MLN8237), exhibits potent toxicity against numerous GBM lines (12-14). Notably, Alisertib can prolong in vivo survival of animals implanted intracranially with glioma cells (15-18). Thus, the drug can cross the blood brain barrier and penetrate intracranial tumor tissue reaching concentrations of up to ∼1000nmol/L (15). The drug also synergizes with TMZ to reduce proliferation in vitro (14). Inhibitors of AURKA, including Alisertib, have reached clinical trials for various cancers including GBM (19, 20), although the drug’s clinical success has not yet delivered the same anti-tumor benefits as observed in preclinical studies.

AURKA controls numerous downstream cellular events (19, 21). One major aspect of the cell cycle regulated by AURKA function is to drive primary cilia disassembly (11, 22-24). Disassembling the primary cilium is an essential step for cells to exit G1 and re-purpose their centrioles for spindle pole formation during G2/M phase. Inhibiting AURKA activity with Alisertib can prevent cilia disassembly and as a result stabilize or increase the frequency of ciliogenesis. For example, Alisertib increased the frequency of cilia in human retinal pigmented epithelial cells (ARPE-19) and human diploid fibroblasts (WI-38) (25, 26). Furthermore, Alisertib restored the frequency of primary cilia in Panc1 cells (a pancreatic ductal adenocarcinoma cell line) (27) and more recently, it restored cilia in supratentorial RELA ependymal cells (28).

To our knowledge, nobody has examined whether or how Alisertib affects GBM cilia, and if it does, whether this treatment could potentially work in concert with TTFields. In this study, we sought to determine whether Alisertib affects the frequency of GBM cilia, and whether these effects could render cells more susceptible to the tumor cell killing effects by TTFields.

## Results

### Inhibiting AURKA via Alisertib reduces the frequency of primary cilia in patient-derived GBM cell lines and biopsies treated ex vivo

We first explored the effect of Alisertib on cilia frequency in cultures of patient-derived gliomaspheres. We grew gliomaspheres from two GBM derived lines (L0, R24-3) and one low grade glioma derived line (S7) using concentrations that approximate physiologic levels measured in brain, tumor and plasma (15). We treated spheres with either vehicle, 1uM or 4uM Alisertib for 24hr, fixed, and immunolabeled cryosections of the spheres for pericentriolar material 1 (PCM1), which concentrates around centrioles and ciliary basal bodies) and ARL13B (which enriches along the primary cilia membrane). We then calculated the number of cilia/sphere area. Strikingly, we observed a marked loss of primary cilia in all 3 lines (**Fig. 1A-C, E-G, I-K**). Quantification of the results confirmed significant reduction of the density of cilia in treated spheres in L0 (**Fig. 1D**), R24-3 (**Fig. 1H**) and S7 cells (**Fig. 1L**). The IC50 of Alisertib in GBM cells has been reported to range from ∼3-225nM depending on whether glioma cells were grown as adherent or spheres (12, 29). Some studies have reported that the Alisertib’s effect on increasing or stabilizing the cilia are observed with as low as 10nM of Alisertib (27). Thus, we also examined whether much lower concentrations (10nM or 100nM) increase GBM ciliogenesis. However, after 24 or 72 of treatment, we observed either lack of an effect or reduced ciliogenesis (**Suppl. Fig 1**). Further, we transfected R24-3 cells with cDNA encoding OFD1:mCherry (which localizes around the basal bodies) and ARL13B^WT^:GFP which enriches in the cilium. We then collected timelapse images over a 24hr period. Compared to vehicle treatment, dissolution of the cilium can be observed in cells treated with 1μM Alisertib (**Fig. 1M-O**).

**Fig. 1.**
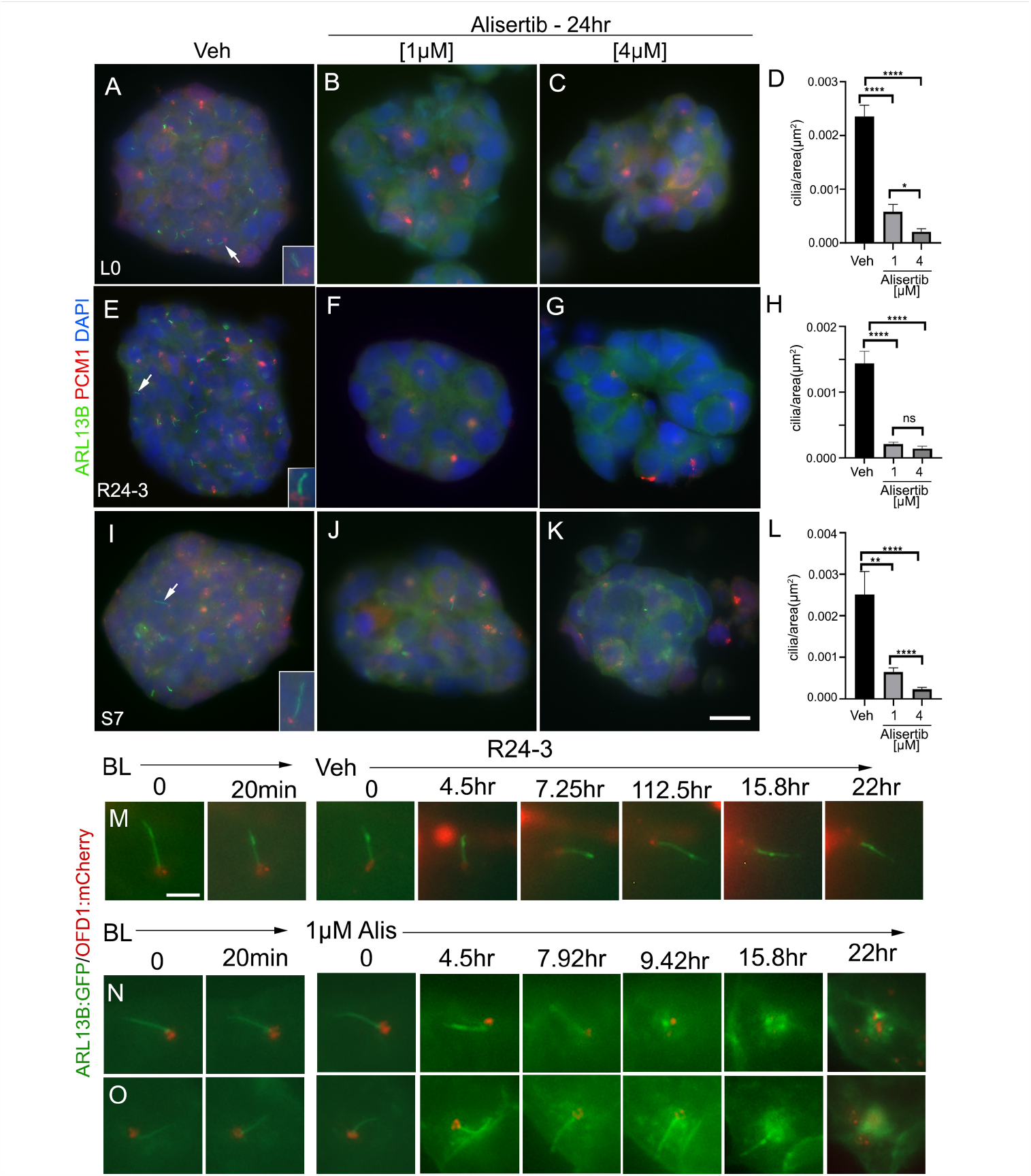
Alisertib lowers frequency of primary cilia in patient-derived GBM spheres in vitro. Patient-derived GBM L0 (A-C), R24-3 (E-G) and low grade gliomaspheres S7 (I-K) exposed to Vehicle or different concentrations of Alisertib. 24hr later, spheres were fixed in PFA, frozen, sectioned and immunolabeled for PCM1 (red), which clusters around basal cilia bodies/centrioles, and ARL13B (green) which enriches in the primary cilium. Nuclei were labeled with DAPI (blue). Arrows and insets (A, E, I) point to cilia in the sphere. D, H, L) The number of ARL13B+ cilia per sphere area in Vehicle or Alisertib-treated cells. ***p<0.001, ****p<0.0001 (ANOVA). M-O) Live imaging of R24-3 cells transiently co-transfected with cDNA encoding Arl13b:EGFP and OFD1:mCherry (which localizes around the base of ARL13B:GFP+ cilia). 24-48hr after transfection cells were timelapsed imaged for about 30min at baseline (BL). Cells were then treated with vehicle (M) or 1μM Alisertib (N, O) and imaged every 10-15 min up to 22 hr later. Note the loss of ARL13B+ cilia up to 22 hr later. Scale bars (μm) in K = 20, M = 5.

We next wondered if Alisertib could have the same influence on cilia in patient biopsies. To that end, d two pathology-confirmed GBM biopsies were subdissected into 3 groups: acute fixation, overnight vehicle, or overnight in 1μM Alisertib (**Fig. 2A**). All groups were fixed, frozen, cryosection and immunolabeled for gamma-tubulin (which labels centrioles/basal bodies) and ARL13B to label the cilium. While we readily observed cilia in tissues that were acutely fixed (**Fig. 2B, E**) and overnight control (**Fig. 2C, F**), cilia frequency was reduced in Alisertib-treated specimens (**Fig. 2D, G**). Quantification of these results showed significant loss of cilia compared to controls in both Alisertib-treated specimens (**Fig. 2H,I**).

**Fig. 2.**
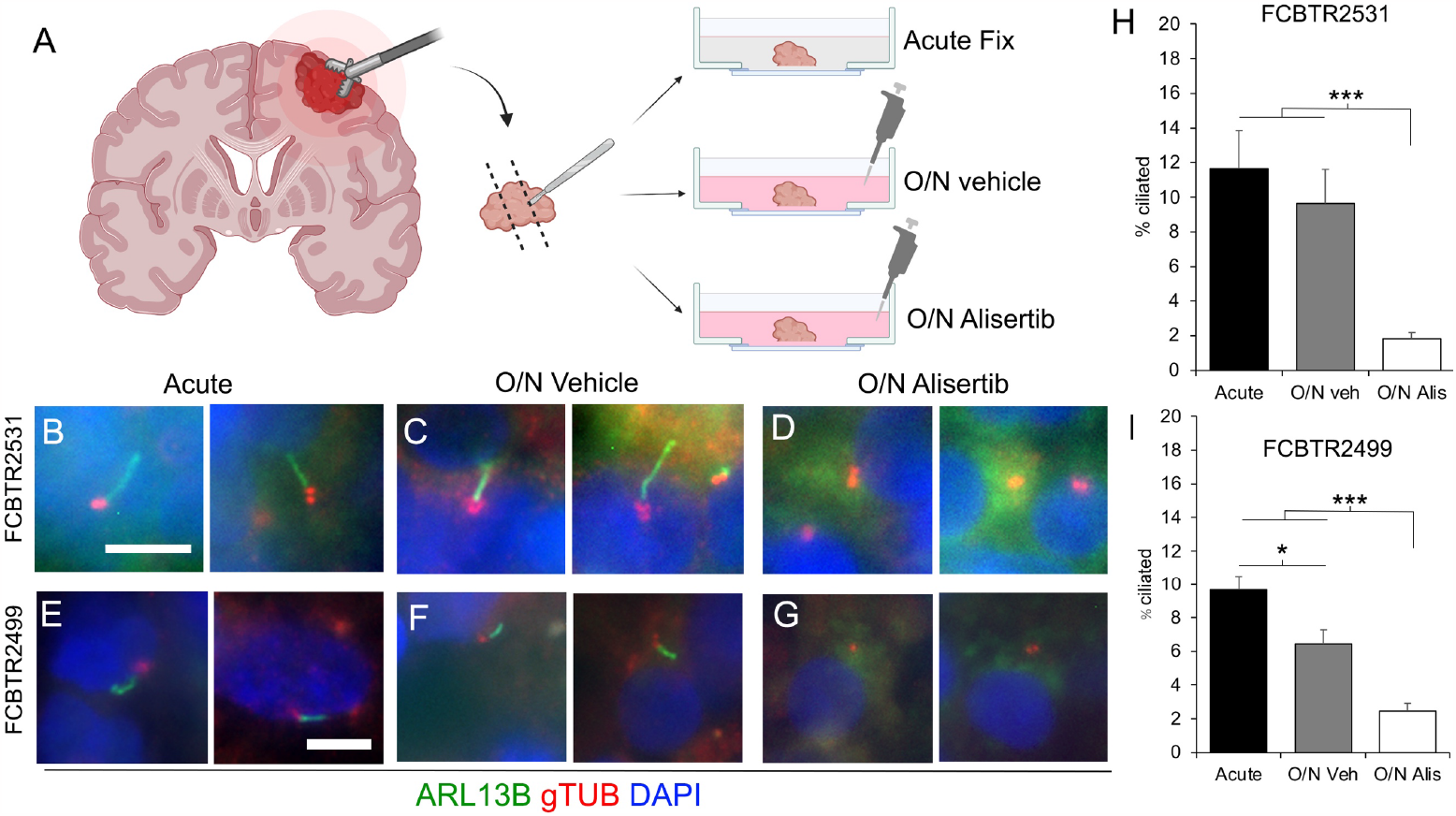
Alisertib disrupts ciliogenesis in GBM patient biopsies treated ex vivo. A) Ex-vivo treatment of surgical resections. Biopsies were dissected and separated into acute fix, overnight (O/N) vehicle or O/N Alisertib (1uM) treatment. B-G) Tissues were fixed, frozen, cryosectioned and immunostained for ARL13B (green) and gTub (red) which labels the ciliary basal body. Nuclei are labeled with DAPI (blue). Two examples of acute fix (B,E), overnight vehicle (C,F) or overnight Alisertib (D,G) from two different patients are shown. H, I) Percent of ciliated cells from indicated patient samples in the indicated group. *p<0.05, ***p<0.001 (ANOVA). Scale bars in B,E = 10μm. Images in A were generated using Biorender.com.

While Alisertib is a selective inhibitor of AURKA, higher concentrations of Alisertib have been reported to inhibit AURKB activity (30). Aurora B/C kinase activity has been linked to glioma cilia disassembly(6). Therefore we wondered if the ciliary changes after Alisertib could be attributable to disruption of AURKB activity in our cell lines. We grew adherent L0 and R24-3 cells in the presence of vehicle or AURKB inhibitor, AZD1152 (Barasertib). Given that as low as 5nM of AZD1152 significantly inhibited GBM cell proliferation in vitro (31), we used a range of 3-100nM of AZD1152 to examine GBM ciliogenesis 24hr after treatment. While we observed changes in nuclear size/morphology/number at all AZD1152 concentrations, we observed no effect of AZD1152 on the percent of ciliated GBM cells (**Suppl. Fig. 2**). These results suggest an AURKA inhibitor-specific effect on glioma cilia depletion.

### Alisertib does not reduce cilia frequency on neurons or glia derived from mouse cortex

Previous studies reported that Alisertib concentrations up to 800nM do not exert appreciable toxicity on normal human astrocytes in vitro (29). Since Alisertib dramatically reduced glioma cilia within 24 hours of exposure, we asked whether cilia of normal primary neural cell types are similarly affected. To test this, we cultured dissociated mouse neonatal cortices on glass coverslips for 12 days in vitro (DIV). At 12DIV, we assigned coverslips as control (vehicle) or 1μM Alisertib. We previously showed that these types of culture allow us to examine the effect of TTFields or drugs on cells that differentiate into various subtypes including astrocytes and neurons, marked by glial fibrillary acidic protein (GFAP) and neuronal nuclei (NeuN) expression respectively (3).

We first examined neuronal cilia by triple immunolabeling for NeuN, type 3 adenylyl cyclase (AC3), an enzyme enriched in most neuronal cilia in the cortex, and pericentrin (Pcnt, a protein concentrated around the cilia basal body) (**Fig. 3A,B**). After 24 hours of Alisertib, we did not observe any significant changes on the frequency of neurons with AC3^+^ cilia (**Fig. 3C**). We next examined astrocyte cilia through a combined immunostaining for GFAP, ARL13B and Pcnt (**Fig. 3D, E**). However after 24 hours of treatment, we did not observe significant differences in the frequency of astrocyte cilia (**Fig. 3F**). These data suggest that, at least at acute timepoints after Alisertib, neuronal and astrocyte cilia are less responsive to Alisertib than glioma cells.

**Fig. 3.**
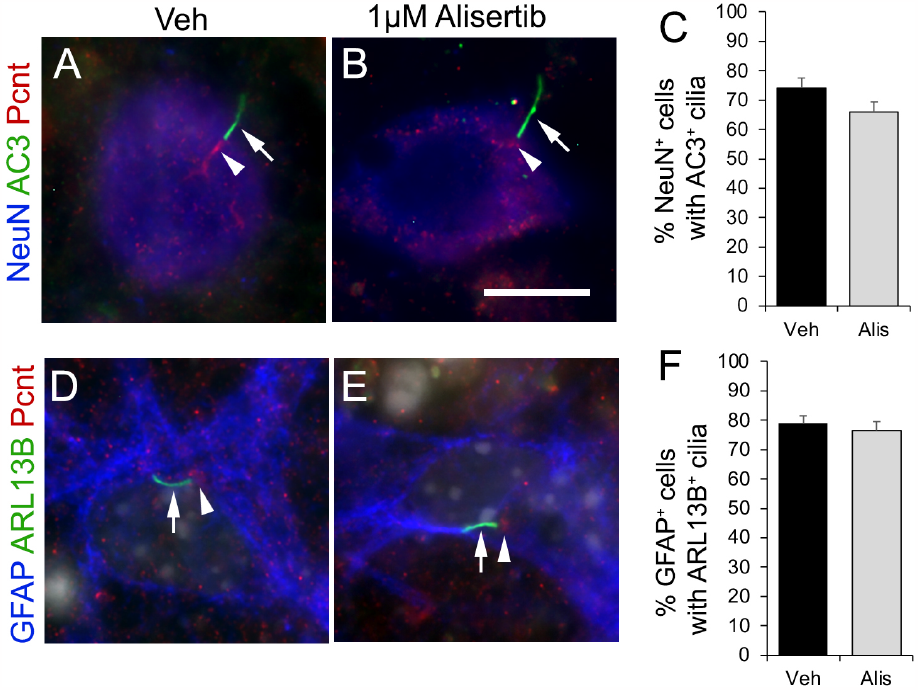
Alisertib does not disrupt neuronal and glial cell primary cilia. Mixed primary cultures from neonatal mouse cerebral cortex were dissociated and maintained for 12DIV and treated with vehicle (Control) or 1μM Alisertib exposed for 24 hour. A,B) Cells were stained for neuronal cilia marker AC3 (green, arrow) and NeuN (blue). Arrows point to AC3^+^ cilia in both groups. C) Percent of NeuN^+^ cells with AC3^+^ cilia. D, E) Cultures were immunostained for ARL13B (green), pericentrin (Pcnt, red), and GFAP (blue). Nuclei are labeled with DAPI (grey). F) Percent of GFAP^+^ cells with ARL13B^+^ cilia. Scale bar in B = 10μm.

### Alisertib with or following TTFields reduces GBM cell proliferation

Our previous study suggested cilia may be associated with tumor cell recurrence following TTFields (3), and that suppressing ciliogenesis may help enhance TTFields efficacy. Since Alisertib reduced GBM cilia frequencies, we wondered if the two treatments may work better together. To test this, we examined L0 and R24-3 cell lines after co-treatment or sequential treatment of TTFields followed by Alisertib (see Fig. 4 top schematics for treatment schedule). The four treatment groups were: vehicle, Alisertib (1μM), vehicle + TTFields, Alisertib + TTFields. We examined proliferation of cells at two points during the treatment schedule. First, we examined the acute total numbers (acute count) of cells immediately after last treatment (i.e. after 1 or 4 days of treatments). Second, we determined the fold expansion (or recurrence index) of cells seven days after the last treatment (fold count). For L0 cells, we found that treating cells with both Alisertib + TTFields significantly reduced cell number (**Fig. 4A,C**) and fold expansion (**Fig. 4B,D**) compared to either treatment alone, and irrespective of the treatment sequence.

**Fig. 4.**
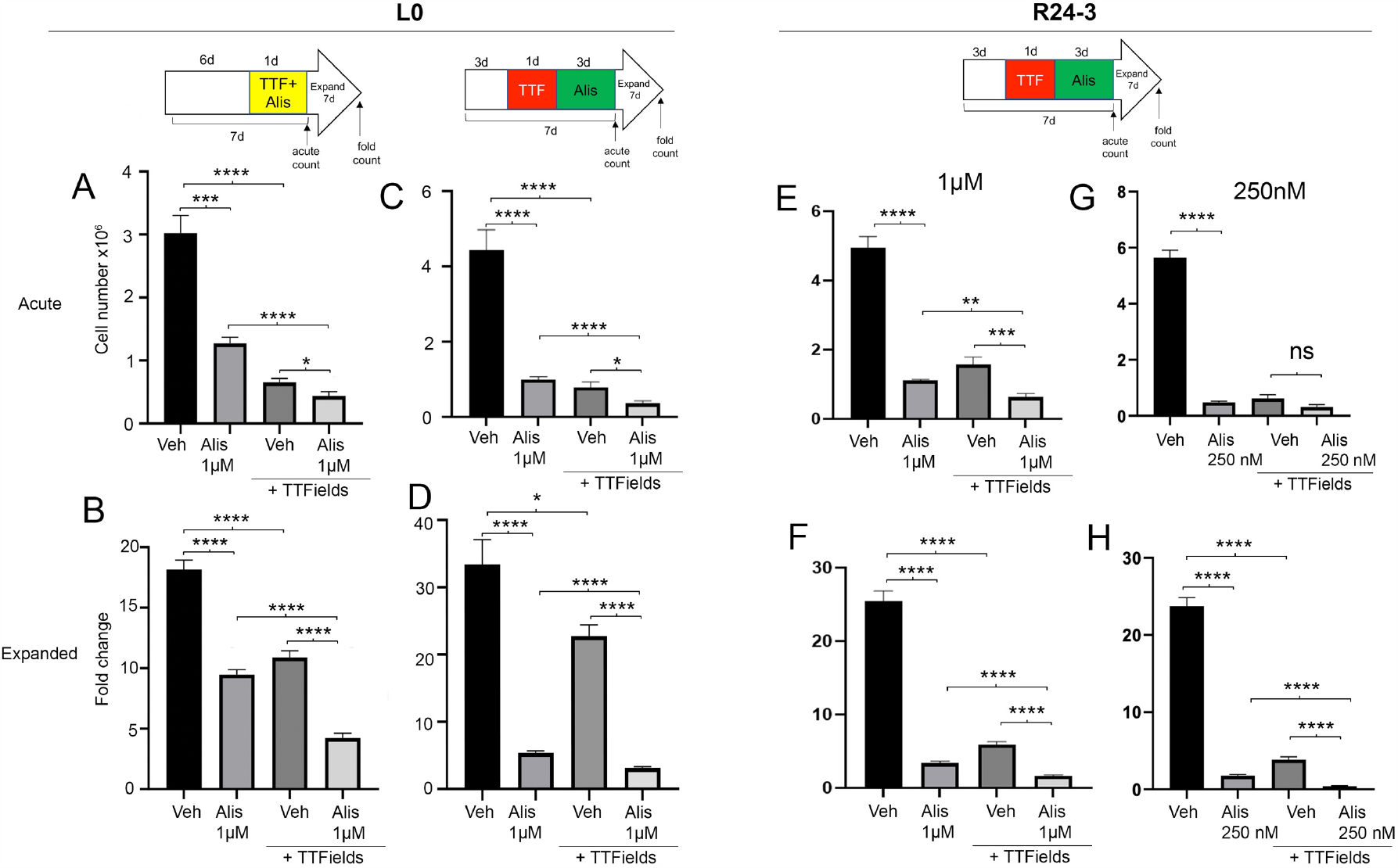
Co- or sequential treatment of TTFields and Alisertib impair GBM cell expansion. L0 (A-D) or R24-3(E-H) cells were grown as spheres for the indicated time and then exposed to either Alisertib alone, TTFields alone or together for 1 day (A,B), or in sequence (C,D, E-H). Cells were counted immediately after treatment at 7 days (Acute count), then pooled, and 2.5x10^4^ cells/well were expanded in fresh media for 7 days in 24 well plates (n=12 wells/group) and the fold expansion was calculated (Fold count). ns =not significant, *p<0.05, **p<0.01, ***p<0.001, ****p<0.0001 (ANOVA)

Similarly in R24-3 cells, Alisertib and TTFields significantly reduced both cell number (**Fig. 4E**) and fold expansion (**Fig. 4F**) of cells compared to either treatment alone. We also tested a lower concentration (250nM) of Alisertib in which we did not find any effect of Alisertib+TTFields at the acute timepoint (**Fig. 4G**), but did observe significantly reduced fold expansion of cells compared to either treatment alone (**Fig. 4H**). We also performed a western blot of treated cell lysates to determine if TTFields itself was contributing to a reduction in phospho-AURKA like 250nM of Alisertib. However we only observed an obvious reduction of phospho-AURKA after Alisertib exposure and not TTFields in both cell lines (**Suppl. Fig. 3**). Altogether, these results indicate adding Alisertib to TTFields treatment has a greater effect of inhibiting GBM cell expansion than either treatment alone.

### Alisertib and TTFields co-treatment effects persist in ARL13B-depleted and ARL13B^+^ cilia-lacking U87-MG GBM cells

We next asked if the effect of Alisertib and TTFields on the GBM cell proliferation in vitro is attributable to Alisertib’s surprising effect on glioma cilia. To test this, we examined two different GBM cell lines lacking cilia: an L0 transgenic cell line that we previously generated by genetic depletion of ARL13B cilia using CRISPR/Cas9 (32), and a U87-MG cell line which are naturally devoid of ARL13B^+^ cilia (33, 34). In L0 cells, we found that Alisertib plus TTFields significantly reduces both acute cell number (**Fig. 5A**) and fold expansion of cells (**Fig. 5B**) compared to either treatment alone. Similarly, in U87-MG cells, we found that the combination of Alisertib and TTFields reduces both acute cell number (**Fig. 5C**) and fold expansion of cells (**Fig. 5D**) compared to either treatment alone. Surprisingly, U87-MG cells appear less sensitive to Alisertib alone (Fig. 5C), though it is not clear if this is associated with the cell’s loss of ARL13B^+^ cilia, presence of serum in U87-MG cells or some other cell-type related factor.

**Fig. 5.**
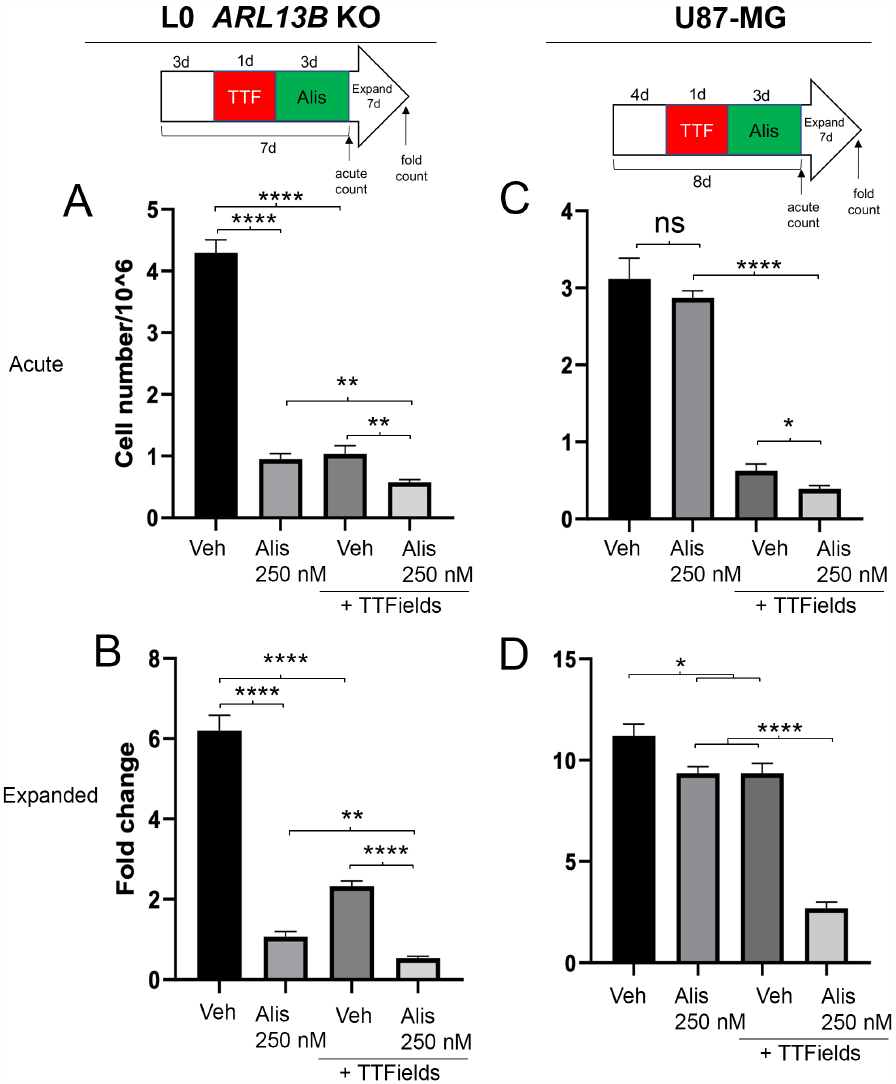
Inhibition of GBM cell proliferation by TTFields and Alisertib is not cilia-dependent. L0 *ARL13B* KO (A,B) or naturally cilia-devoid U87-MG (C,D) cells were grown as spheres (L0) or adherent (U87-MG) for the indicated treatment schedule. Cells were counted immediately after treatment at 8 days (Acute count), then pooled, and 2.5x10^4^ cells/well were expanded in fresh media for 7 days in 24 well plates (n=12 wells/group) and the fold expansion was calculated (Fold count). ns =not significant, *p<0.05, **p<0.01, ***p<0.001, ****p<0.0001 (ANOVA)

Nevertheless, these results suggest Alisertib’s effects on cilia are not required to generate a combined treatment effect observed in wildtype GBM cells.

## Discussion

Our data show that concomitant treatment of Alisertib and TTFields in vitro suppresses GBM cell proliferation greater than either treatment alone. This effect is not mediated via tumor primary cilia, as both deciliated and ciliated populations of GBM cells appear sensitive to the combination of treatments. To our knowledge, we are the first to show that inhibiting AURKA suppresses GBM ciliogenesis both in vitro and ex vivo using patient derived biopsies, an effect not observed in normal mouse cortical neurons or glia. The unexpected cilia ablation may serve as a useful readout of the drug’s penetrance into GBM tumors, a possibility that requires further in vivo investigation.

### Significance of Alisertib effects on glioma primary cilia

Considering that AURKA plays a key role in ciliary disassembly, and GBM cells constantly assemble and disassemble their cilia while proliferating, it was surprising to not observe increased GBM cilia frequency after Alisertib treatment. It is possible that AURKA inhibition in GBM cells stimulates activity of other molecules that promote ciliary disassembly such as HDAC6, NEK and others (for review see: (22, 35, 36)). However, we previously reported that overexpression of HDAC6 in our L0 or S7 glioma cells did not reduce cilia frequency or length (32). Alisertib may have a selective capacity to kill GBM cells in their ciliated state. In part this may explain why U87-MG cells were less sensitive to Alisertib alone treatment (Fig. 5C), as U87-MG cells naturally lack cilia (34). In contrast, we found that L0 GBM cells lacking cilia were highly sensitive to Alisertib (Fig. 5a), which could be attributable to secondary downstream effects of CRISPR/Cas9-mediated depletion of ARL13B. We recently showed that TTFields-induced loss of primary cilia is in part attributable to activation of autophagy (3). Considering Alisertib also induces autophagy in human glioblastoma cells (13), future studies should examine whether Alisertib-induced autophagy is mediating GBM cilia depletion. Lee and colleagues recently found that inhibiting Aurora B/C kinases using GSK-1070916 increased the frequency of ciliated patient-derived glioma stem cells (6). However, when we inhibited AURKB specifically with AZD1152, we saw no effect on the frequency of ciliated GBM cells. The difference between the two studies could be related to use of a different drug, different cell lines, or an unappreciated role of Aurora C kinase in ciliogenesis.

Whatever the mechanism associated with AURKA-induced depletion of cilia, the presence of cilia was not required for combined benefit of TTFields and Alisertib to suppress proliferation.

The observation that Alisertib does not depend on primary cilia to work in concert with TTFields is not surprising. Like TTFields (for review see: (2)), AURKA regulates many other cellular processes with pro-tumorigenic activity (for review see: (19)) which their inhibition by Alisertib can potentially be enhanced by concomitant application of TTFields. This raises the question of whether the ciliary disruptions we observe are relevant. First, because Alisertib suppressed ciliogenesis in cultured biopsied tissue (Fig. 2) and the drug is measurable in CSF, brain and intracranial tumors, the drug-induced loss of tumor cilia in vivo could be a potential biomarker of Alisertib activity in the tumor. Second, tumor cilia are induced by and promote resistance to standard of care TMZ treatment in vitro and in vivo (3, 8-10). Considering Alisertib plus TMZ has been reported more toxic during gliomagenesis than either treatment alone (14), future studies should examine how Alisertib and TTFields, which suppress ciliogenesis, interact with TMZ chemotherapy which appears to stimulate ciliogenesis.

### Is the AURKA pathway a viable adjunct therapy with TTFields?

Since TTFields prolong patient survival, there are active searches for combinatorial drug targets, particularly those that interfere with tumor cell mitosis and cell survival mechanisms. Relevant to our study, an inhibitor of Aurora B kinase pathway, AZD1152 (Barasertib) displays both properties by disrupting mitosis and survival of newly diagnosed and recurrent GBM cells in vitro which is exacerbated by TTFields treatment (31). Our in vitro data suggest similar effects may be achievable by targeting Aurora A kinase pathway, a widely studied target across cancers (19, 37). Future in vivo studies should compare Alisertib or other AURKA inhibitors with TTFields.

Although Alisertib has reached Phase 1 clinical trials for high grade glioma (20), it appears that their results have not yielded the same success as seen in preclinical studies. Some of the challenges in working with Alisertib could be related to drug-induced changes to tumor cells that may countereffect anti-tumor immunity, or that effective dosages required to reach/kill brain tumors are significantly lower than needed. For example, a recent study suggests Alisertib may induce PD-L1 expression in tumor cells, which serves to suppress anti-tumor immunity (38). However in our hands, Alisertib did not have an appreciable effect on GBM cell PD-L1 expression by western blot (data not shown). Further, while Alisertib has been shown by multiple groups to reach the brain and slow brain tumor growth in vivo as described above, a recent study examining Alisertib penetrance into CNS suggests its distribution and concentrations are limited (39). Supporting this observation, Alisertib treatment alone did not slow intracranial tumor growth of transplanted GL261glioma cells, but did synergize with co-treatment of Birabresib (a BET bromodomain inhibitor) to significantly extend survival(40). Thus it is feasible that low doses of Alisertib in the brain could combine or potentiate other drug or brain targeted therapies such as TTFields. Taken together our results demonstrate a potential benefit for concomitant application of TTFields and Alisertib and warrants further investigation in vivo possibly with concurrent use of other GBM clinically relevant drugs.

## Materials and Methods

### Cell culture

Human L0 (grade IV glioblastoma from a 43 year old male) and S7(grade II glioma from a 54 year old female) cell lines were isolated and maintained as previously described (41-44). R24-03 cell lines were expanded from a 91year old male GBM . Human U87-MG (Cat# HTB-14) cells were obtained from ATCC (Gaithersburg, MD, USA) and maintained in recommended media conditions. L0, R24-3, and S7 cells were grown as floating spheres and maintained in NeuroCult NS-A Proliferation medium and 10% proliferation supplement (STEMCELL Technologies, Vancouver, BC, Canada; Cat# 05750 and #05753), 1% penicillin–streptomycin (Thermofisher, Waltham, MA, USA; Cat# 15140122), 20 ng/ml human epidermal growth factor (hEGF) (Cat #78006), and 10 ng/ml basic fibroblast growth factor (bFGF) (STEMCELL Technologies; Cat #78003). For S7 cells, the media was supplemented with 2 μg/ml heparin (STEMCELL Technologies; Cat #07980). When cells reached confluency, or spheres reached approximately 150 μm in diameter, they were enzymatically dissociated by digestion with Accumax (Innovative Cell Technologies, San Diego, CA, USA; Cat#AM-105) for 10 min at 37 °C. For human cells grown on glass coverslips, NeuroCult NS-A Proliferation medium was supplemented with 10% heat inactivated fetal bovine serum (FBS) (Cytiva, Marlborough, MA, USA; Cat #SH30070.03HI). All cells were grown in a humidified incubator at 37 °C with 5% CO_2_. Where indicated, indicated cell lines were treated with Alisertib (Selleckchem.com; Houston, TX, USA; Cat# S1133), or AZD1152 (MedChemExpress, Monmouth Junction, NJ USA; Cat # HY-10127) which were dissolved in 100% DMSO (Fisher Scientific; Cat # D128-500).and fixed after indicated treatment durations and processed as described below.

Primary neural cultures were obtained and grown as previously described (3). Briefly, acutely micro-dissected C57/BL6 mouse cortices from postnatal day 0-2 pups were dissected into Gey’s Balanced Salt Solution (Sigma-Aldrich, Cat #G9779) at ∼37 °C under oxygenation for ∼20 min. Dissociated cells were triturated with pipettes of decreasing bore size, pelleted by centrifugation at 1,500 rpm for 3-5 min, and resuspended and plated in glial medium containing DMEM (Cytiva HyClone, Cat# SH3002201), FBS (Gemini BioProducts, West Sacramento, CA, USA; Cat# 50-753-2981), insulin (Sigma-Aldrich, Cat# 15500), Glutamax (Gibco, Waltham, MA, USA;Cat# 35050061) and Penicillin-streptomycin (Gibco, Cat# 15140122). Cells were plated at a density of 80,000 cells/coverslip on 12-mm glass coverslips coated with 0.1 mg/ml poly-D-lysine followed by 5 μg/ml laminin in minimal essential medium. After approximately 2 hrs, cells were supplemented with 2mL neuronal media containing Neurobasal A (Gibco, Cat# 10888022) supplemented with B27 (Gibco, Cat# A3582801), Glutamax (Gibco, 35050061), Kynurenic acid (Sigma Aldrich, Cat# K3375), and GDNF (Sigma Aldrich, Cat# SRP3200 ). Every 4 days, half of the media was replaced with fresh neuronal media as described above but lacking kynurenic acid and GDNF. On DIV12, coverslips were treated with vehicle or Alisertib and fixed after 24 hrs.

### Ex-vivo culture and Alisertib treatment

In accordance with our institutional IRB protocol (#201902489), we collected several fresh, surgically-resected tumor biopsies that were subsequently pathologically confirmed. Within 1 hour of the resection, biopsies were taken to the laboratory, and dissected into several pieces using a sterile scalpel blade. Tissues were immediately fixed and/or transferred into 2mL of S7 media for culture at 37°C in 5% CO_2_ and treated with vehicle or Alisertib, and fixed 24hr later in 4% paraformaldehyde.

### TTFields application

Adherent cells (U87-MG) or spheres (L0, R24-3) were placed in TTFields ceramic dishes, each dish approximately 25mm in diameter, and mounted onto inovitro™ base plates (Novocure Ltd., Haifa, Israel). The base plates were connected to a power generator which delivered TTFields at frequency of 200 kHz at a target intensity of 1.62V/cm (45). Treatment duration are as indicated, ranging from 24 to 72 hours for a single treatment. To prevent media evaporation during TTFields application, parafilm was placed over each TTFields ceramic dish. Control samples were grown in 6 well plates. Data in each TTFields experiment were technical replicates and pooled from at least 6-8 dishes per condition and per timepoint.

### Cell Proliferation Assessment

We examined proliferation of cells at two points during our treatment schedules. First, we calculated acute numbers of cells immediately after last treatment (i.e., after 1 or 4 days of treatment(s)). Total cell counts from each condition were collected using a Bio-Rad TC20 automated cell counter (Bio-Rad, Hercules, CA, USA). Second, we examined the fold expansion (or recurrence index) of cells seven days after the last treatment by seeding cells (2.5x10^4^) in 500 μl of growth media per well in 24-well plates for each experimental group (n=12 wells per group). After 7 days, cells were enzymatically dissociated and resuspended in 1X phosphate-buffered saline (PBS) and counted. Mean fold changes +/- SEM were plotted and compared to vehicle-treated controls. Data were analyzed statistically using one-way ANOVA using Prism (v9.5.1).

### Timelapse imaging

For timelase imaging, we plated 50,000 cells in R24-3 media supplemented with 5% FBS into 35 mm glass bottom culture dishes (Ibidi, Gråfelfing, Germany; cat #81158) which were maintained at 37°C in 5% CO_2_. Twenty four hours before imaging at about 70% confluency, cells were transfected with 500 ng total cDNA/dish of pDest-Arl13b:GFP (a kind gift from T. Caspary, Emory University) and pCMV-myc/mCherry:hOFD1 (Vectorbuilder.com, vector ID: VB201119-1128fyp) using Lipofectamine 3000 (Life Technologies; Carlsbad, CA, USA; cat#L3000015). Imaging was conducted on an inverted Zeiss AxioObserver D1 microscope using a Zeiss 40×/0.95 plan Apochromat air objective. The microscope stage was equipped with a Tokai Hit stage incubation system that maintained a humid environment and stage temperature of 37°C and 5% CO_2_. After approximate 30 minutes of baseline images were obtained, vehicle/drug treatment was applied and images were acquired every 5 minutes for up to 24hours. Exposure times ranged in duration from 400 to 750msec (EGFP) and 300-400 msec (Cy3) per image. Image acquisition and processing were performed using Zeiss ZEN software (ZEN 2012 (Blue edition) v1.1.2.0).

### Western Blot

Western blot was performed as recently described (34). Briefly, cells were harvested at indicated time points and lysed in 1x cell lysis buffer (Cell Signaling Technologies, Inc, Danvers, MA, USA; Cat#9803) or 1× radioimmunoprecipitation assay (RIPA) buffer (Cell Signaling; Cat# 501015489) containing 1× protease inhibitor cocktail (Sigma, St. Louis, MO, USA; Cat# P2850), phosphatase inhibitor cocktails 1 (Sigma, St. Louis, MO, USA; Cat# P5726), and 2 (Sigma; Cat# P0044), and 1× phenylmethanesulfonyl fluoride (Sigma; Cat# 93482). Prior to loading, samples were heated to 70°C for 10 min. A total of 25-30 μg of total protein lysate per lane were separated on 4–12% Bis-Tris gels (Thermofisher; Cat# NP0050). Proteins were blotted onto PVDF membranes using iBlot (program 3 for 8 min; Invitrogen, Carlsbad, CA, USA). Blots were blocked in 5% nonfat dry milk (NFDM) or bovine serum albumin (BSA, Jackson Immuno Research, West Grove, PA, USA; Cat# NC9871802) in 1× tris-buffered saline (TBS) with 0.1% Tween (TBST) for 20 min and then incubated in primary antibodies in 2.5% NFDM or BSA in 1× TBST for 24 h at 4 °C. Blots were rinsed and probed in the appropriate horseradish peroxidase (HRP)-conjugated secondary antibody (1:10,000; BioRad, Hercules, CA, USA) for 30 min at RT in 2.5% NFDM or BSA in 1× TBST. Finally, blots were rinsed in 1× TBS and developed using an Amersham ECL chemiluminescence kit (Global Life Sciences Solutions USA, Marlborough, MA, USA), and images were captured using an AlphaInnotech Fluorchem Q Imaging System (Protein Simple, San Jose, CA, USA). Selected areas surrounding the predicted molecular weight of the protein of interest were extracted from whole blot images.

### Immunostaining

For immunocytochemical (ICC) and immunohistochemical (IHC) analyses, samples were fixed at indicated timepoints with 4% paraformaldehyde in 0.1 M phosphate buffer (4% PFA) for 30min for 15minutes (ICC) to 1 hour (IHC) and washed with 1x PBS. Spheres or biopsies were cryoprotected in 30% sucrose in PBS followed by a 1:1: 30% sucrose and optimal cutting temperature compound (OCT) (Fisher Healthcare, #4585), frozen in OCT over liquid N2 and cryosectioned at 16μm. Samples were stained for the indicated primary antibodies: mouse anti-beta-actin (1:10,000; Sigma; Cat #A5316), mouse anti-gamma Tub (1:3,000; Sigma; Cat# T6557), mouse anti-ARL13B (1:3000; Abcam, Waltham, MA USA; Cat# AB136648), rabbit anti-ARL13B (1:3000; Proteintech, Rosemont, IL USA; Cat # 17711-1-AP), chicken anti-GFAP (1:1000; Encor Biotechnology, Gainesville, FL USA; Cat# CPCA-GFAP), chicken anti-GFP (1:5000; Abcam; #ab13970), rabbit anti-pericentrin (1:1000; Covance; Cat #: PRB-432C), chicken anti-type 3 adenylyl cyclase (1:5000; Encor Biotechnology; Cat# CPCA-ACIII), and rabbit anti-PCM1 (1:1000; Bethyl Laboratories, Montgomery, TX; Cat# A301-150A). Samples were incubated in blocking solution containing 5% normal donkey serum (NDS) (Jackson Immunoresearch, West Grove, PA, USA; Cat#NC9624464) and 0.2% Triton-X 100 in 1x PBS for 1 hour and then incubated in primary antibodies with 2.5% NDS and 0.1% Triton-X 100 in 1x PBS either for 2 hours at room temperature (RT) or overnight at 4°C. Appropriate FITC-, Cy3- or Cy5-conjugated secondary antibodies (1:1000; Jackson ImmunoResearch) in 2.5% NDS with 1x PBS were applied for 1-2 hour at RT, and coverslips were mounted onto Superfrost™ Plus coated glass slides (Fisher Scientific, cat # 12-550-15) in Prolong Gold antifade media containing DAPI (Thermofisher; Cat# P36935). Stained coverslips were examined under epifluorescence using an inverted Zeiss AxioObserver D1 microscope using a Zeiss 40×/0.95 plan Apochromat air objective or a Zeiss 63X/1.4 plan Apochromat oil objective. Images were captured and analyzed using Zeiss ZEN software.

## Acknowledgements

Funding for this research was supported by a 2022 AACR-Novocure Tumor Treating Fields Research Grant (#22-60-62-SARK) (to. M.R.S).

---

**Suppl Fig. 1.**
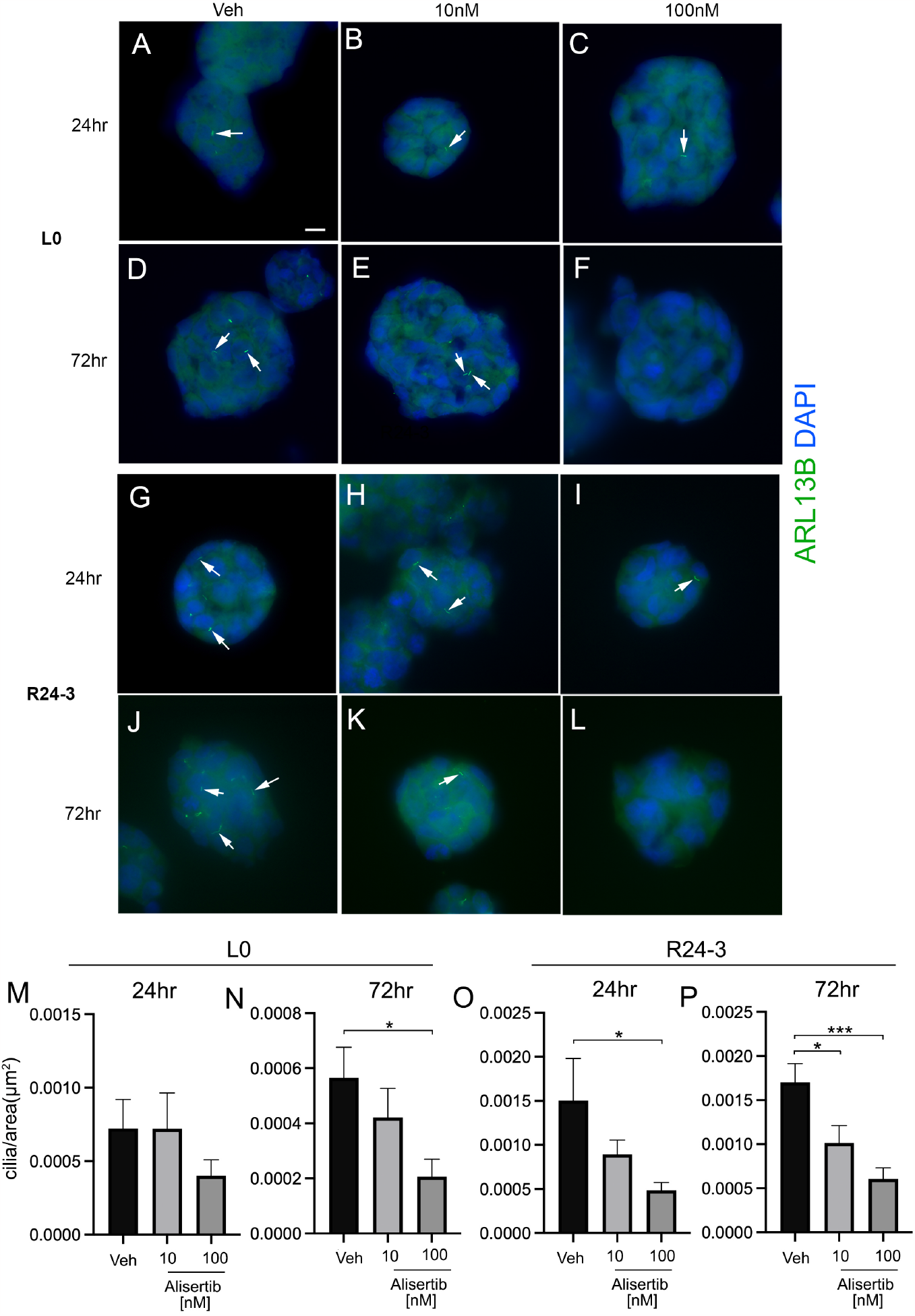
Low concentrations of Alisertib do not affect or significantly reduce GBM cilia frequency. L0 and R24-3 spheres were treated with vehicle, 10nM or 100nM Alisertib. Spheres were fixed after 24 or 72hr, frozen sectioned and immunostained for ARL13B. A-L) Examples of spheres from each treatment group. Arrows point to ARL13B+ cilia, nuclei are labeled with DAPI. Scale bar in A = 10μm. M-P) Quantification of the# of cilia/area for indicate cell line and treatment group. *p<0.05, ***p<0.001 (ANOVA).

**Suppl Fig. 2.**
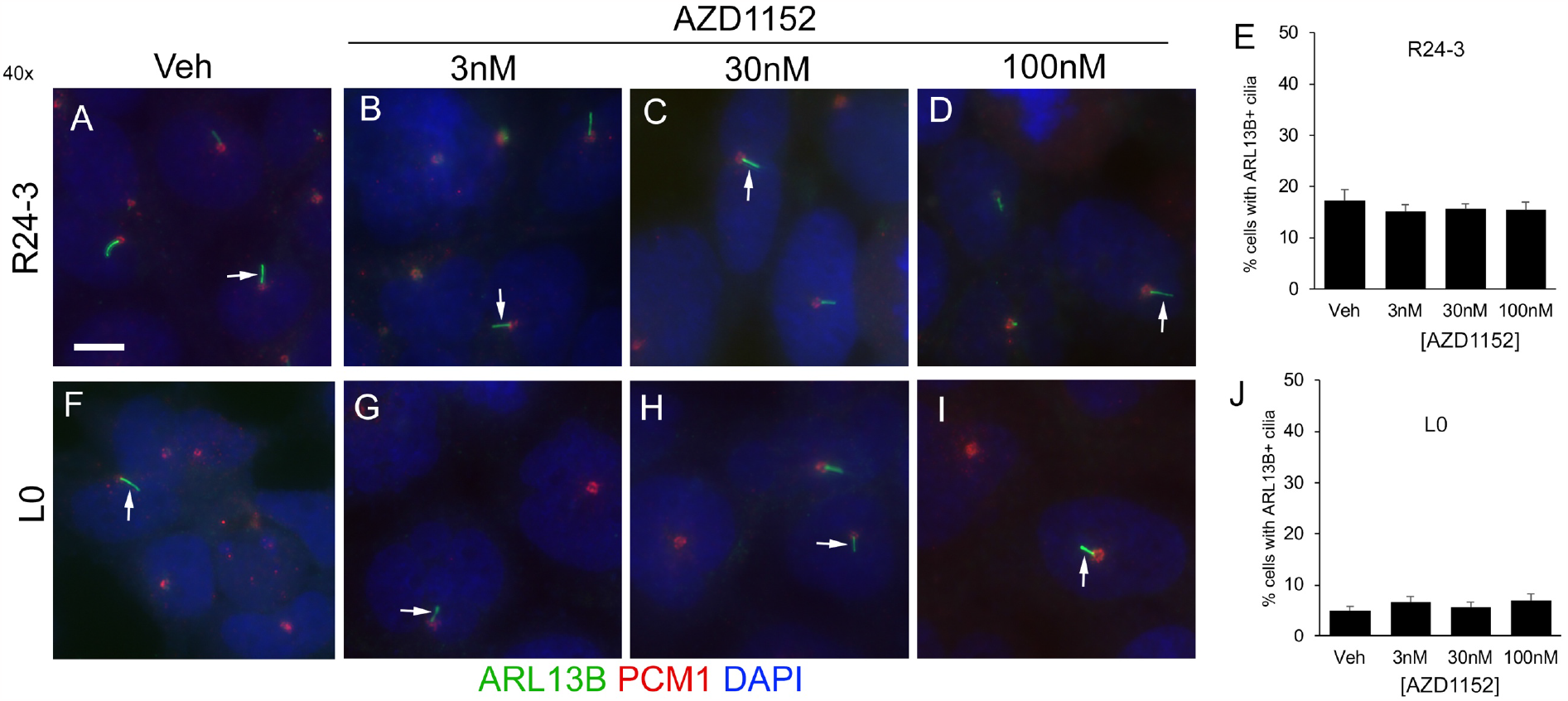
AURKB inhibitor, AZD1152 does not affect GBM ciliogenesis. Adherent R24-3 (A-D) and L0 (F-I) cells were treated with vehicle, 3nM, 30nM or 100nM AZD1152. Cells were fixed after 24hr and immunostained for ARL13B (green) and PCM1 (red), and counterlabeled with DAPI (blue). Arrows point to ARL13B^+^ cilia. Scale bar in A = 10μm. E and J) Percent of cells with ARL13B^+^ cilia for R24-3 (E) and L0 (J) cells for the indicated treatment/concentration.

**Suppl Fig. 3.**
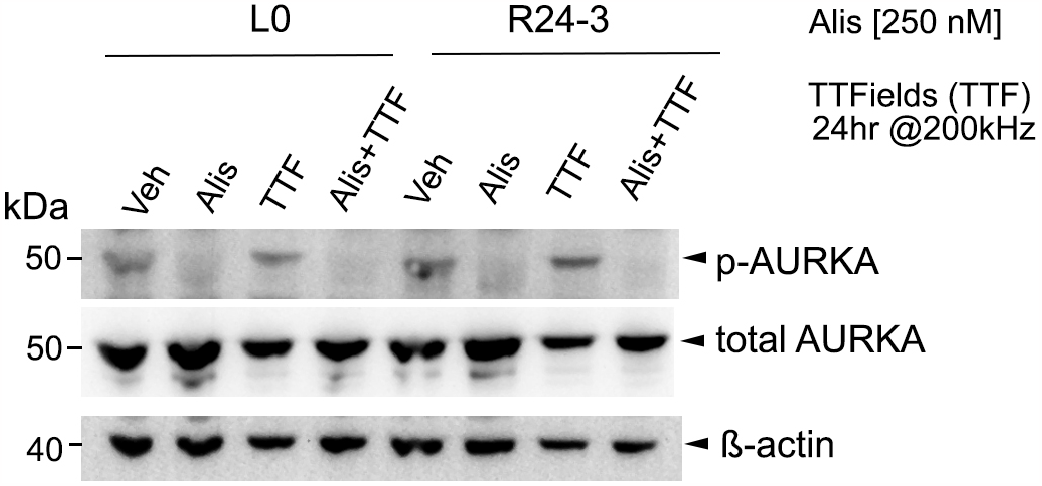
Alisertib, but not TTFields, reduces p-AURKA protein in GBM cell lines. L0 and R24-3 spheres were treated with vehicle (Veh), 250nM Alisertib (Alis), TTFields alone, or Alis and TTFields. After 24hr, spheres were lysed and protein lysates generated. Western blots were probed for p-AURKA, stripped and re-probed for total AURKA and ß-actin.

